# Brain wave dynamics in Hopfield Kuramoto model

**DOI:** 10.1101/2024.08.13.607707

**Authors:** Ruwei Yao, Yichao Li, Xintong Yao, Kang Wang, Jingling Qu, Bo Hong, Xiaolong Zou

## Abstract

Whole brain neural oscillation activities exhibit multiple wave patterns and seem to be supported by the common circuit network structure. We proposed a Hebbian-like Kuramoto model based entirely on heterogeneous connectivity strength rather than phase delay, which encodes the multiple phase patterns as attractors. We systematically investigated how the model dynamic landscape influenced by attractors and their corresponding eigenvalues, as well as how to control the stability of equilibrium points and the occurrence of high dimension bifurcations. This framework enables us to reproduce the dominant wave activity components in human brain functional MRI signal, and provides a canonical model for the multi body physical system spatio-temporal pattern attractor dynamics.

## I. INTRODUCTION

Multi-body physical systems can be abstracted using complex network structures characterized by connectivity matrices [1]. These systems often exhibit intricate spatiotemporal activity patterns [2, 3]. Research in neuroscience has shown that neural system activity can manifest as regular and robust spatiotemporal activity patterns. Advanced and sophisticated methods have been developed to extract and analyze these neural patterns from raw neural data [4, 5]. However at the same time, the underlying relationship between network circuit and these activity patterns remains under discussion. For this topic, exploring a canonical framework, particularly through the computational modeling approaches such as the Hopfield models [6, 7] and the continuous attractor neural networks [8], has also been widely called for [9, 10].

Recent discoveries in human brain EEG [11, 12], ECoG [13, 14] and fMRI [15, 16] have identified regular spatiotemporal phase patterns, known as the whole brain large-scale waves phenomenon. Large-scale waves, including modes such as traveling and standing waves, have been shown to be closely related to cognitive function [15, 17]. Numerous studies in computational neuroscience have aimed to explain the generation mechanism of these wave phase patterns through whole brain dynamic modeling [18]. By establishing the differential equation of oscillators or wave equation, combined with the white matter fibers connection extracted from the Diffusion Tensor Imaging (DTI) [19, 20] or cortex geometry information [21, 22], the whole brain model can simultaneously exhibit a range of phase patterns, including plane and rotating waves.

While the aforementioned whole brain modeling driven by structural foundations has achieved success, there remain several details that need further exploration. The First is the mechanism behind the specific spatial phase patterns of large-scale brain waves. Research indicates that both the connection transmission delays and the connectivity strengths can influence the phase relationships between nodes [23]. However, the theoretical determination of factors, particularly connectivity strength, in generative modeling of specific traveling waves still necessitates innovative research. Another important question is, in the case of coexisting patterns, how the model network parameters influence the dominance and stability of wave patterns, as well as the global energy landscape [24]. Answering these questions will provide a theoretical framework for modeling neural wave activities and understanding their structural basis in the brain.

In this paper, we sought to leverage the classic Kuramoto model [25, 26] to capture the attractor dynamics in the Hopfield network, thereby simulating multiple brain wave phase patterns. We first derived the characteristic equation satisfied by the equilibrium-point phase-locked manifold based on the mean field method [25, 26]. Subsequently, we proposed a Hebbian-like approach for encoding multiple wave patterns into the recurrent connectivity matrix of the Kuramoto model. In contrast to previous Kuramoto wave generation models dependent on phase delays [27, 28] or models for binary patterns [29], we proposed a method entirely based on heterogeneous connectivity strength, validating its capacity for memorizing multiple wave phase attractors. We further explored the influence of memorized phase patterns and eigenvalues of the connectivity matrix on shaping the model potential function. This exploration enabled us to attain control over the dominance and stability of various patterns, as well as the generation of high-dimensional bifurcations. Our theoretical framework establishes a canonical form for a brain wave generation model, distinct from previous simulation models with given structural network. This framework is positioned to inspire novel approaches to understanding wave activity through the lens of brain dynamic landscapes.

## II. KURAMOTO PHASE LOCKED

We first considered the original Kuramoto model with *L* coupled oscillators defined by Eq. (1).

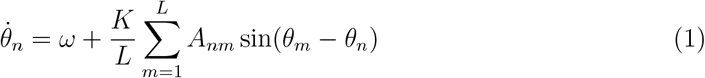

Where *θ*_*n*_ ∈ (−*π, π*] denotes the phase of node *n, ω* denotes the intrinsic angular frequency, *K* is the global coupling strength and *A*_*nm*_ ∈ ℝ represents the effective connectivity strength from node *m* to node *n*. While the Kuramoto model is typically used for the study of synchronization or phase transition [26], in this context, we primarily focused on its ability to generate specific wave patterns. For this aim we introduced two related concepts. For a *L* coupled oscillators dynamic system, suppose ***θ***(*t*) = {*θ*_*n*_(*t*)}_*L*×1_ is the solution of Eq. (1), define that the system Eq. (1) reaches the phase locked state if for each 1 ≤ *n, m* ≤ *L, n* ≠ *m*, there is

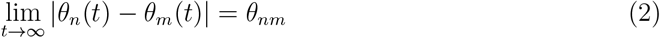

Define that the system Eq. (1) reaches the frequency synchronization state if for each 1 ≤ *n, m* ≤ *L, n* ≠ *m*, there is

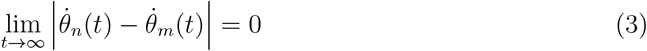

Consider a simple kind of phase-locked manifold that all oscillators eventually oscillate at their intrinsic frequency *ω*, which means ∀*n*, 1 ≤ *n* ≤ *L*

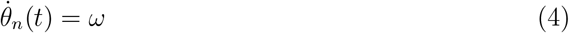

Such manifold satisfies phase locked definition in Eq. (2) and frequency synchronization definition in Eq. (3). We have derived the necessary condition for the existence of such phase-locked manifold in Eq. (4). Defining the phase bias of each node *ψ*_*n*_ = *θ*_*n*_ − *ωt*, it could yield the differential equation Eq. (5) of the phase bias *ψ*_*n*_.

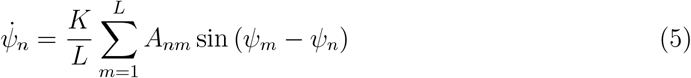

For system Eq. (5), we used the complex mean field method to derive the characteristic equation satisfied by the equilibrium points 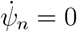. (see Appendix A for details).

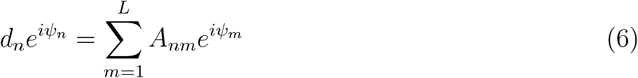

Where ***ψ*** = {*ψ*_*n*_}_*L*×1_ ∈ 𝕋^*L*^ is the solution of Eq. (1), *d*_*n*_ ∈ R, *d*_*n*_ *>* 0, 1 ≤ *n* ≤ *L*, and the matrix form

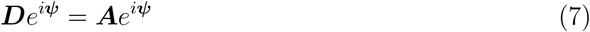

Where ***D*** = diag (*d*_1_, *d*_2_, …, *d*_*L*_). We named Eq. (6, 7) the phase pattern characteristic equation. It is also the necessary condition for the existence of a phase-locked manifold Eq. (4) on the *L*-torus 𝕋^*L*^. From Eq. (7) we can see that the phase-locked manifold ***ψ*** is a generalized complex eigenvector of the connectivity matrix ***A*** = (*A*_*nm*_)_*L*×*L*_. Additionally, one should note that multiplying both sides of the Eq. (7) by *e*^*iωt*^ does not change the equation, which ensures the circumferential invariance of the phase-locked manifold.

Given a phase pattern that meeting the phase pattern characteristic equation, it could be high dimensional attractors (stable) or high dimensional saddle points (unstable). To find the stable phase-locked manifold, we calculated the Jacobian matrix ***J***(***ψ***) = {*J*_*nm*_(***ψ***)}_*L*×*L*_ of the system Eq. (5) at the equilibrium point in Eq. (8) and verify its local linearization stability (see Appendix B for details). The real parts of the Jacobian matrix eigenvalues provides information on local invariant stability, instability, and central manifold of equilibrium points.

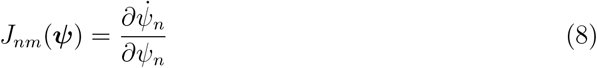

To quantitatively characterize the Kuramoto model dynamics landscape and the current state of the system, we defined the scalar function *P*(***ψ***(*t*)) of the phase state variables.

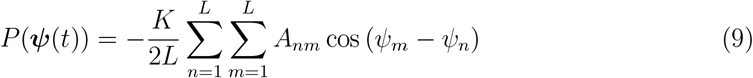

Where ***ψ***(*t*) = {*ψ*_*n*_(*t*)}_*L*×1_ is solution the of Eq. (5). We employed the function *P*(***ψ***(*t*)) as a descriptor of the dynamical landscape of the system Eq. (1) or Eq. (5) on 𝕋^*L*^. To illustrate, consider a common scenario where the matrix ***A*** is a real symmetric matrix [29], it is easy to verify that *P* (***ψ***(*t*)) satisfies

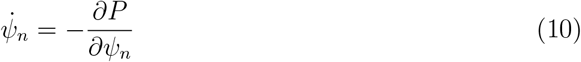

Considering the change of the *P*(***ψ***(*t*)) over time, we yield

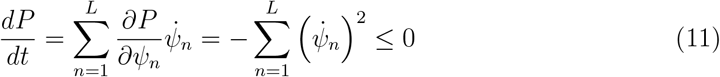

Eq. (11) shows that the scalar function *P*(***ψ***(*t*)) only decreases with the evolution of the dynamic system. In this case, *P*(***ψ***(*t*)) is the potential function, known as the energy function at times, of the gradient system Eq. (5). In addition, *P*(***ψ***(*t*)) is a bounded function and is not constant within the domain of definition if ***A*** ≠ ***O***, thus ensuring the asymptotic stability of the equilibrium point.

## III. CONNECTIVITY STRENGTH BASED METHOD

Here we aimed to modify the Kuramoto model to simultaneously exhibit specific distinct target wave patterns. A feasible approach is the multiple attractors scheme. This approach involves constructing a specific connectivity matrix that allows multiple wave phase patterns to become the stable phase-locked manifolds of the model. Subsequently, starting from the random initial phase pattern, the system will probabilistically converge into different wave phase pattern attractors and stabilize in a specific phase-locked state. This process is reminiscent of the classical Hopfield network [6], which can memorize specific binary patterns, While the binary patterns serve as the discrete attractors of the Hopfield network, the phase pattern complex vectors on the unit circle are the attractors of Kuramoto model after our construction.

As we mentioned in the last section, wave phase pattern attractors must satisfy the phase pattern characteristic equation. Hence we can use the Hebb’s rule of association of the classical Hopfield network for reference. For memorize a phase patterns 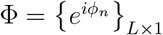, considering the Hebb outer product sum method, we have

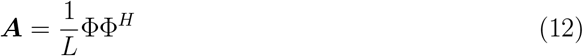

So there are 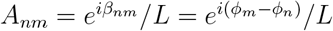, we can verify that ***A***Φ = Φ. Next, combining Eq. (A2), We have

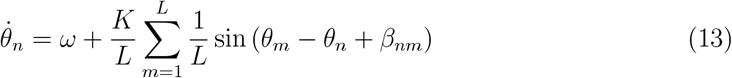

Where *β*_*nm*_ is the phase delay between node *n* and *m*. The above process is the classic Hebbain kuramoto method [27, 28]. It can be seen that the Hebb outer product sum method in the complex field is equivalent to a system with added phase delay, which memorizes phase patterns dependent on the phase delay mechanism. We refer to this approach, as described in Eq. (13), as the connectivity delay based method.

Here, we proposed a method for constructing the connectivity matrix that does not require any explicit phase delay. Instead, it relies entirely on the heterogeneous connectivity strength between nodes. Suppose that we need to memorize *S* phase patterns Φ_1_, Φ_2_, …, Φ_*s*_ ∈ ℂ^*L*×1^, where 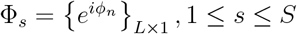. Let ***Φ***_*s*_ ∈ ℝ^*L*×2^ be a matrix formed by concatenating the real and imaginary parts of Φ_*s*_.

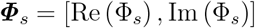

And let **Φ** = [***Φ***_1_, …, ***Φ***_*S*_] ∈ ℝ^*L*×2*S*^. Considering the special case of the phase pattern characteristic equation, that is for phase pattern ***Φ***_*s*_, We let ***D*** = *λ*_*s*_***I***_*L*×*L*_ where *λ*_*s*_ ∈ ℝ, so we yield

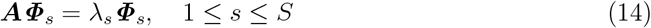

Thus obtaining a solution for the connectivity matrix ***A***

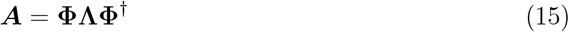

Where **Φ**^†^ = (**Φ**^*T*^ **Φ)**^−1^ **Φ**^*T*^ denotes the pseudo inverse matrix of **Φ**, and **Λ** = diag (*λ*_1_, *λ*_1_, …, *λ*_*S*_, *λ*_*S*_) is the diagonal matrix whose diagonal elements are the corresponding eigenvalues of ***A***. The rank of ***A*** is 2*S*. A more general solution family of connectivity matrices can be found in the Appendix C. It is straightforward to verify that the matrix ***A*** defined by Eq. (15) satisfies the phase characteristic equation Eq. (7). In the subsequent sections, we will further elucidate the significance of the eigenvalue matrix **Λ** to the model and how to select the eigenvalues *λ*_*s*_ to make its corresponding Φ_*s*_ become a stable phase-locked manifolds. Additionally, It can be observed that the connectivity matrix ***A*** in Eq. (15) is a pure real matrix and therefore does not explicitly contain a delay term. We refer to Eq. (15) as the connectivity strength based method.

## IV. RESULT

We first studied the performance of our method when memorizing a single wave phase pattern. We conducted simulation of the Kuramoto system Eq. (1) on the 6 × 6 lattice with *L* = 36 nodes. We designed a toy target phase pattern 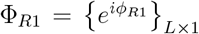, which rotates one revolution counterclockwise on the lattice [Fig. 1(a)]. In actual whole brain modeling, the target phase patterns is usually obtained through the phase analysis of the recorded electrophysiological data. Here, we hope that the model can stably converge to the target phase pattern. We constructed model connectivity matrix using connectivity strength based method and connectivity delay based method, respectively. Then we initialized ***θ***(*t* = 0) ∈ ℝ^*L*^ by sampling ***θ***(*t* = 0) of each node from a uniform distribution between [0, 2*π*), and use the same initial value for two times simulations. We set the intrinsic angular frequency *ω* = 10 × 2*π*, the global coupling strength *K* = 200, set the simulation step to 0.01 s and simulate 10 s (1000 timepoints) of Eq. (4). For the connectivity strength based method, Fig. 1(b) shows the constructed connectivity matrix. Generally speaking, the smaller the phase difference between *ϕ*_*n*_ and *ϕ*_*m*_ is, the larger *A*_*nm*_ will be, which is in line with Hebbian rule’ s principle “Fire together, wire together”. We further verified that the model converges to the target phase pattern. We extracted the phase pattern complex vector *e*^*i**θ***(*t*)^ ∈ ℂ^*L*^ at *t* = 10*s* and verified that *e*^*i**θ***(*t*=10)^ and ***A*** indeed satisfies the phase pattern characteristic equation [Fig. 1(b)].

**FIG. 1.**
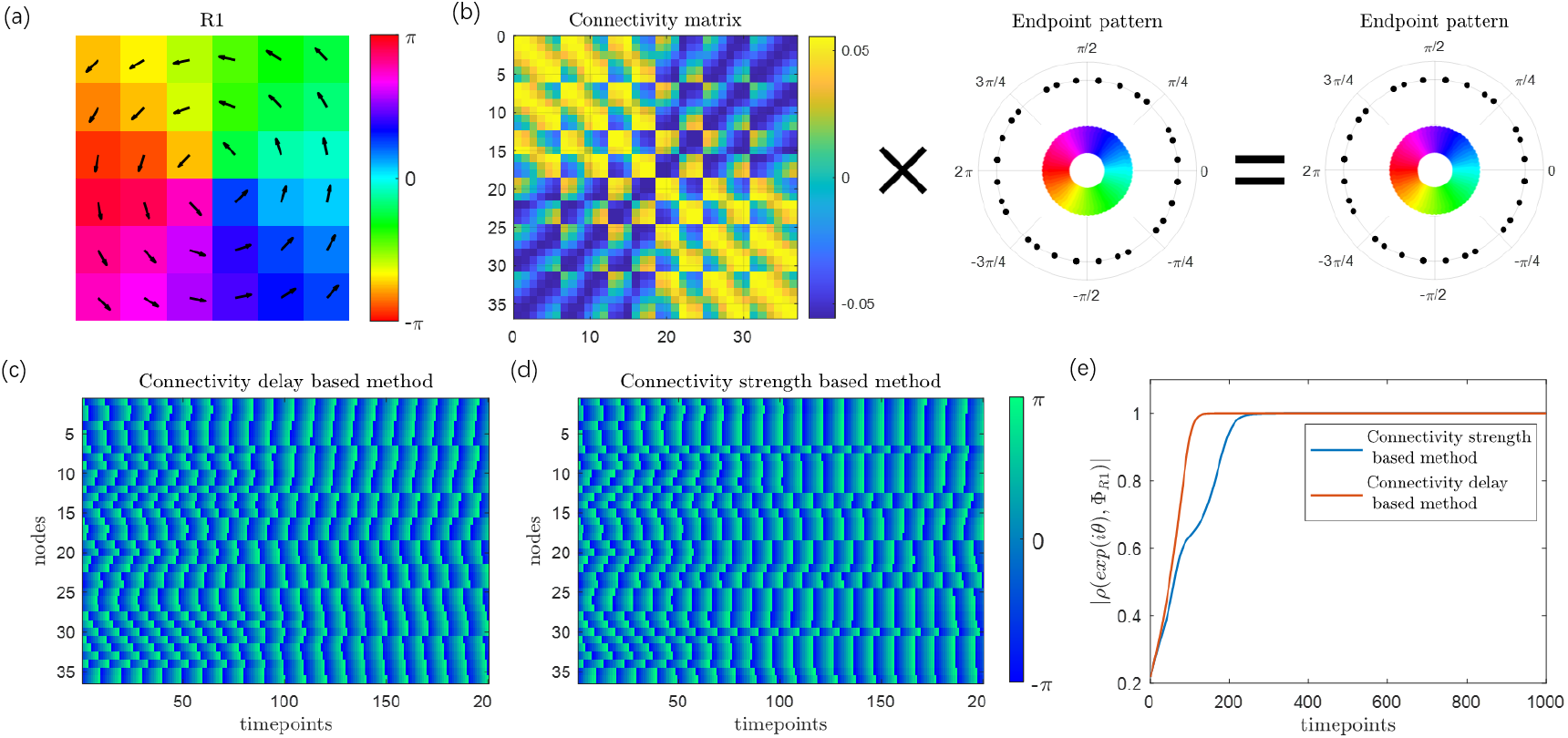
The connectivity matrix construction methods memorize specific traveling wave phase patterns. (a) The memorized phase pattern is a traveling wave that rotates counterclockwise on a two-dimensional lattice, with the winding number 1. The 6 × 6 lattice’s central coordinates is [0, 0]. The arrows indicate the phase gradient field, and the direction of the traveling wave phase gradient field on each grid is perpendicular to the line connecting the grid and the center. (b) The connectivity matrix constructed by the Connectivity strength based method. The left multiplication of the connectivity matrix and the phase pattern at the 10s simulation obtains an identity map. (c-d) The simulation phase diagram of (c) Connectivity strength based method and (d) Connectivity delay based method within 2s. (e) The absolute value of affinity function within 1000 time points (10s). The red line represents the Connectivity delay based method, the blue line represents the Connectivity strength based method.

At the same time, we compared the characteristics of the connectivity strength based method Fig. 1(c) and the connectivity delay based method Fig. 1(d). The phase heatmap of the first 200 time points (2s) shows the difference in phase state evolution between them. To quantitatively reflect this time process, we calculated the affinity between the simulated phase patterns {*e*^*i**θ***(*t*)^}_*L*×1_ and the memorized phase pattern Φ_*R*1_. We defined the Affinity function between the two phase patterns 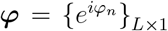 and 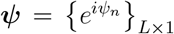 as their normalized inner product Eq. (16)

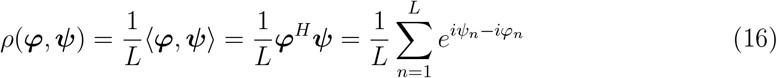

So there is |*ρ*(***φ, ψ***)| = 1 when ***φ, ψ*** is completely equal and |*ρ*(***φ, ψ***)| ≈ 0 when they are completely different. The curve of affinity function show that both the two methods can let the model converge to the memorized phase pattern, but with different trajectories and velocities [Fig. 1(e), red and blue curve].

It should be noted that since the connectivity strength based method get the real number connectivity matrix, when any phase pattern Φ_*s*_ is stored, its conjugate phase pattern 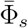 will have the completely equal potential value and Jacobian matrix with Φ_*s*_, and thus has the completely equal stability, which means that Φ_*s*_ and 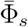 will be memorized at the same time. This characteristic has been observed in classic Hopfield networks.

Next, we attempt to memorize multiple wave phase patterns by connectivity strength based method, and study how our method shapes the global dynamical landscape. We memorized two pairs of conjugate phase patterns on the same lattice structure Fig. 2(a) using the connectivity strength based method in Eq. (15). The first pair is Φ_*R*1_ and its conjugate phase pattern clockwise rotational pattern Φ_*R*2_. The second pair is a phase pattern that translate from top to bottom Φ_*D*1_ (the phase of the lowest row is 5*π*/6, the phase step between adjacent rows is *π*/6) and its conjugate translational pattern Φ_*D*2_ from bottom to top. While the above phase patterns in our demonstration constitute specific instances, rotational and translational waves are prevalent solution forms in numerous Reaction-Diffusion Equations [30, 31], as their spatial structure just in accordance with the low frequency components of spatial harmonics on the lattice. The specific construction process of ***A*** is visually depicted in Fig. 2(b), where *λ*_*R*_ are the eigenvalues corresponding to the rotational pattern and *λ*_*D*_ are the eigenvalues corresponding to the translational pattern. We always constrained *λ*_*R*_ + *λ*_*D*_ = 1 for normalization in subsequent discussions. Eq. (15) ensures that the two pairs of phase patterns are equilibrium points, and the stability of the equilibrium points will be discussed in the next section.

**FIG. 2.**
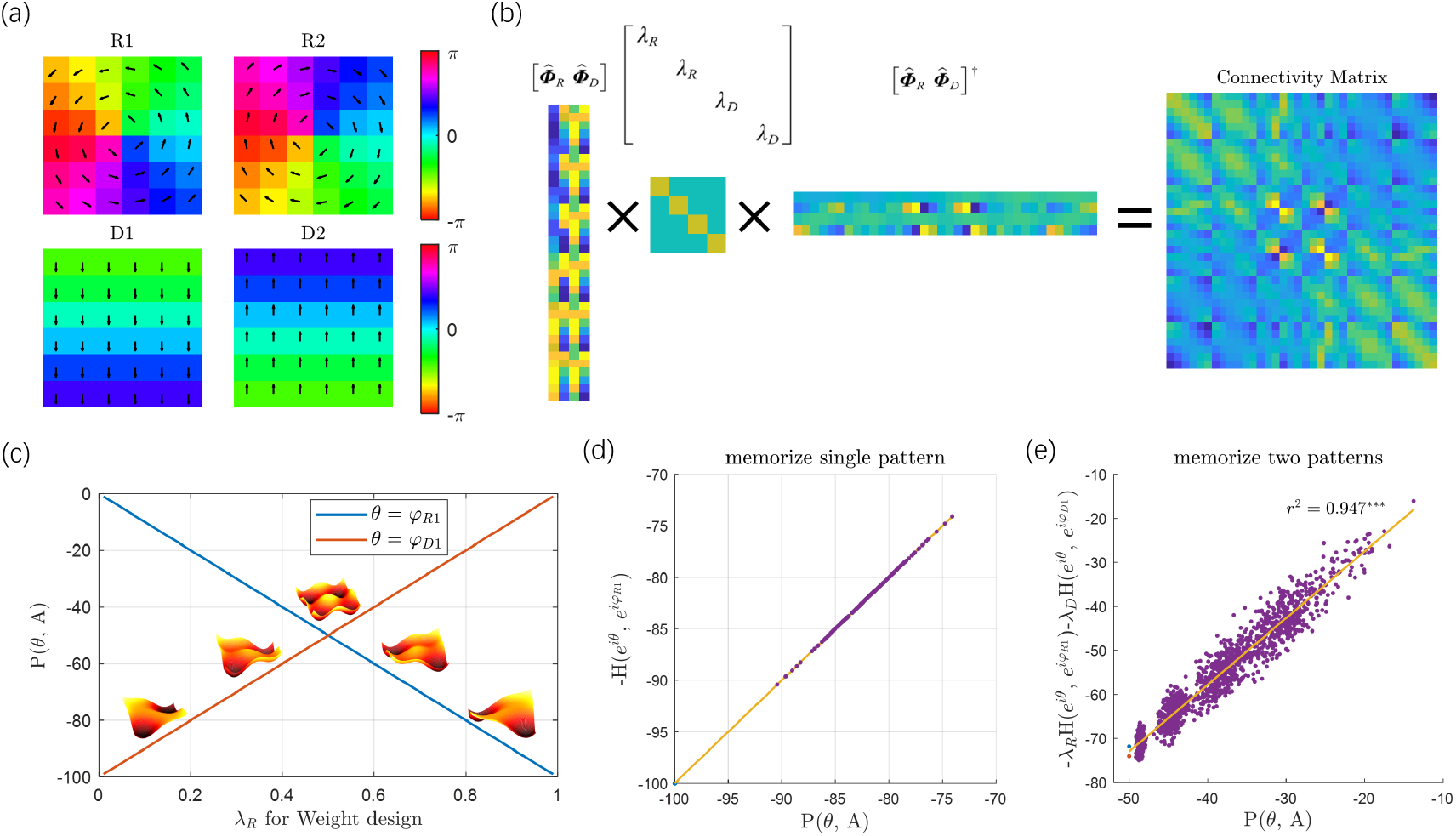
Dynamical landscape properties when memorize multiple phase patterns. (a) Memorize two pairs of conjugate phase patterns: counterclockwise(R1) and clockwise(R2) rotational traveling waves with winding number 1, and translational traveling waves that shift from top to bottom(D1) and from bottom to top(D2). Arrows indicates the phase gradient field. (b) Connectivity strength based method with the eigenvalue configuration *λ*_*R*_ = 0.5, *λ*_*D*_ = 0.5. (c) The relationship between the *P*(*θ*) function value of the target phase patterns and the selected *λ*_*R*_ (maintained *λ*_*R*_ +*λ*_*D*_ = 1) for the connectivity matrix, with blue line indicating the rotational pattern and red line indicating the translational pattern. Surface diagram represents the cartoon dynamical landscape. (d-e) The relationship between potential function value and the affinity with attractors when memorizing (d) single phase pattern Φ_*R*_ or memorizing (e) multiple phase patterns Φ_*R*_ and Φ_*D*_, *λ*_*R*_ = 0.5, *λ*_*D*_ = 0.5. Each purple dot represents a randomly sampled phase pattern, the blue dot represents the standard rotational phase pattern, the red dot represents the standard translational phase pattern, and the yellow line represents the fitting line (*r*^2^ = 0.947 in (e)).

We first examined the relationship between the eigenvalues of the connectivity matrix *λ*_*R*_, *λ*_*D*_ we selected and an overview of the dynamical landscape at Φ_*R*_ and Φ_*D*_. We uniformly increased the *λ*_*R*_ from 0.01 to 0.99 to parameterize the connectivity matrix by eigenvalues and calculate the function *P*(***θ***) values for Φ_*R*_ and Φ_*D*_. The result [Fig. 2 (c)] shows that the function *P*(***θ***) value of the target phase patterns Φ_*R*_ (Φ_*D*_) decreases linearly with the increase of its corresponding eigenvalues *λ*_*R*_ (*λ*_*D*_). It means that the larger the eigenvalues assigned to a target phase patterns, the deeper the phase patterns will be in the dynamical landscape (see Fig. 2(c) cartoon figure), and its stability will be stronger compared to other memorized phase patterns. This provides a guidance for us to adjust the weight of different wave attractors.

Based on the above results, we further studied how the wave phase patterns attractors affect the model dynamical landscape. Attractors are local minimums of the potential function. An intuitive experience is that the closer a phase patterns is to the attractor, the lower its potential value. As for the classic Hopfield Network, the energy function can be represented by the overlay between the current state and the memorized patterns [32]. Therefore, we explored the phase affinity function. Considering the equivalence of the conjugate phase patterns, we defined the Square Affinity Function(SAF) Eq. (17) used to measure the closeness of any phase patterns *e*^*i**θ***^ to the memorized phase pattern pair Φ and 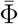.

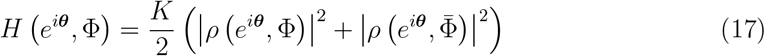

In Appendix D, we proved that when memorizing a single target phase pattern that satisfies certain conditions through the connectivity strength based method, the potential function in Eq. (9) is equal to the opposite number of SAF in Eq. (17). Fig. 2(d) shows 200 random phase pattern samples for verification.

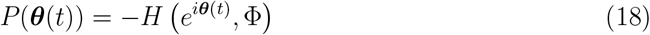

Where the symmetric connectivity matrix ***A*** = **ΦΦ**^†^. Here we demonstrated that when multiple phase patterns with imperfect orthogonality are memorized, the potential function shares a similar relationship with SAF as well, and the SAF for each memorized phase pattern is weighted by their corresponding eigenvalues. To illustrate this, we randomly sampled 2000 real number phase vectors (Add Gaussian noise with zero mean, 0.2, 0.4, 0.6, 0.8, 1.0 five variance levels to the two memorized phase patterns, with each reference phase patterns sampled 200 times at each noise level) for verification. We chose *λ*_*R*_ = 0.50, *λ*_*D*_ = 0.50 for model and calculated the potential function values of the samples under the current model, while also calculated the SAF of the samples with standard rotational and translational patterns respectively. Linear regression analysis was conducted on the potential function and eigenvalue weighted SAF Fig. 2(e). The results show that there is also a high linear correlation (regression *r*^2^ = 0.947) for memorizing multiple phase patterns. The above property clearly reveals the relationship between the dynamical landscape and attractor patterns as well as their eigenvalues.

So far, we have not systematically analyzed the stability of phase patterns equilibrium points memorized by connectivity strength based methods. The stability of these equilibrium points in dynamic systems determines whether the system’s trajectories will stably converge to the equilibrium state over a sufficiently long period of time, which is crucial for understanding the steady states of brain dynamics. In nonlinear systems, the stability of equilibrium points is often dependent on model parameters. When an equilibrium point transitions from stable to unstable as a result of parameter changes, this transition is known as a bifurcation. Our research explores the high-dimensional bifurcation phenomena that arise when multiple phase patterns are memorized using our method. We identified the eigenvalues associated with the target phase patterns as critical parameters for this analysis.

We uniformly increased *λ*_*R*_ from 0.01 to 0.99 and calculate a family of parameterized connectivity matrices according to Eq. (15). For the model with each connectivity matrix, we calculated the Jacobian matrix at the equilibrium point phase pattern Φ_*R*_ and Φ_*D*_ according to Eq. (B1), and calculate the eigenvalues of the Jacobian matrix. We found that these eigenvalues are all real numbers, which is determined by our construction method. Then we sorted these Jacobian matrix eigenvalues from largest to smallest, and focus on the maximum eigenvalues. Fig. 3(a) shows the change of Jacobian matrix eigenvalues at Φ_*R*_ and Φ_*D*_ with parameters *λ*_*R*_. For Φ_*R*_, its Jacobian matrix eigenvalues greater and less than 0 at the same time in the interval [0, 0.394), which means it is a high dimension saddle point. With the increase of *λ*_*R*_, all the Jacobian matrix eigenvalues gradually decrease piecewise linearly. When *λ*_*R*_ = 0.394, the maximum Jacobian matrix eigenvalue is no longer greater than 0, and a high dimension saddle-node bifurcation occurs. With the increase of *λ*_*R*_, the maximum Jacobian matrix eigenvalue maintain 0, Φ_*R*_ remains as the stable attractor. The zero eigenvalue of Jacobian matrix indicates the center manifold of a continuous attractor, corresponding to the circumference invariance of Φ_*R*_. For Φ_*D*_, in the interval (0.539, 1], its Jacobian matrix has eigenvalues greater and less than 0 and Φ_*D*_ is a high dimension saddle point. The saddle-node bifurcation occurs at *λ*_*R*_ = 0.539, and then with the increase (decrease) of *λ*_*D*_ (*λ*_*R*_) it remains a stable attractor.

**FIG. 3.**
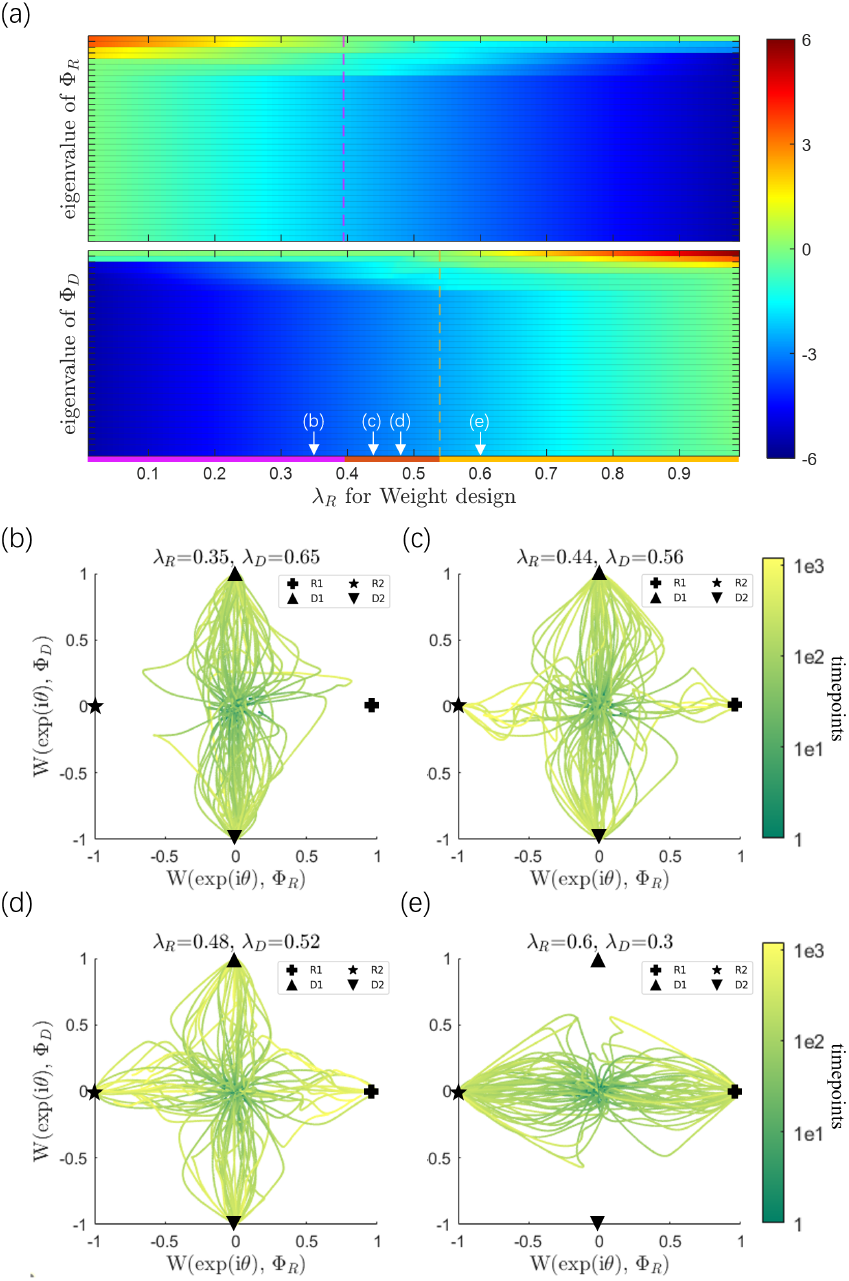
The changes of eigenvalues cause high dimensional bifurcations in the model. (a) The Jacobian matrix eigenvalues at the equilibrium point phase patterns indicate the bifurcations with the change of parameters *λ*_*R*_. The horizontal axis is configured *λ*_*R*_ (maintained *λ*_*R*_ + *λ*_*D*_ = 1) for the connectivity matrix. The upper (lower) part of the diagram shows the 36 eigenvalues (all real numbers, arranged from top to bottom according to the largest to smallest) of the Jacobian matrix at the equilibrium point Φ_*R*_ (Φ_*D*_). The pink (orange) line in the upper (lower) diagram indicates the boundary where the maximum Jacobian matrix eigenvalue at Φ_*R*_ (Φ_*D*_) is equal to 0. The pink interval in the bottom line indicates Φ_*D*_ stable and Φ_*R*_ unstable, the red interval indicates Φ_*R*_ and Φ_*D*_ are both stable, and the orange interval indicates Φ_*R*_ stable and Φ_*D*_ unstable. The white arrows indicate the parameter values corresponding to the subplots b-e. (b-e) 100 times of random initial value simulation in 1200 timepoints. Phase patterns fall into different attractors with different parameters (b) *λ*_*R*_ = 0.35, *λ*_*D*_ = 0.65 (c) *λ*_*R*_ = 0.44, *λ*_*D*_ = 0.56 (d) *λ*_*R*_ = 0.48, *λ*_*D*_ = 0.52 (e) *λ*_*R*_ = 0.6, *λ*_*D*_ = 0.3. The colorbar indicates simulation time point.

In order to demonstrate the stability changes of equilibrium point phase patterns caused by the bifurcation, we selected *λ*_*R*_ in different intervals to construct connectivity matrices for simulation. Maintaining the identical model parameter settings and phase initialization scheme as mentioned above, we conducted 100 rounds of random initial phase simulations for each connectivity matrix. For visualization, we projected the high dimensional simulated phase patterns of first 1200 timepoints onto a two dimensional plane [Fig. 3(b,e)]. With the aspiration that this projection may effectively reveal the similarity between any phase patterns and the attractors, we defined the W function *W* (*e*^*i**θ***^, Φ) between two phase patterns *e*^*i**θ***^ and Φ based on the affinity.

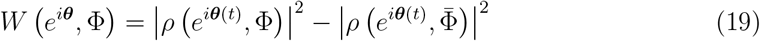

When *e*^*i**θ***^ and Φ are equal there is *W* (*e*^*i**θ***^, Φ) = 1, when *e*^*i**θ***^ and 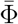 are equal there is *W* (*e*^*i**θ***^, Φ) = −1. If *e*^*i**θ***^ is completely different from Φ or 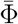 there is *W* (*e*^*i**θ***^, Φ) ≈ 0.

For any phase pattern, we set the horizontal axis as *W* (*e*^*i**θ***^, Φ_*R*1_) and the vertical axis as *W* (*e*^*i**θ***^, Φ_*D*1_) to project the *e*^*i**θ***^. On this plane, the two pairs of standard traveling wave patterns in Fig. 2(a) as the reference fall on the midpoint of the four sides [Fig. 3(b,e)], and the randomly initialized phases are basically projected onto the center position of the plane. We observed that when *λ*_*R*_ is small Fig. 3(b), the translational pattern is the attractor while the rotational pattern is not, resulting in all phase patterns converging to Φ_*D*1_ or Φ_*D*2_. When *λ*_*R*_ = 0.44, *λ*_*D*_ = 0.56 both Φ_*R*_ and Φ_*D*_ are attractors Fig. 3(c), the phase patterns begins to converge to the rotational pattern with a certain probability, and when *λ*_*R*_ = 0.48, *λ*_*D*_ = 0.52 this probability increases Fig. 3(d), it indicates the change of the respective domains of attraction of Φ_*R*_ and Φ_*D*_. With the *λ*_*R*_ increase, the concaves on the landscape corresponding to the rotational pattern deepens and its attraction domain becomes wider, gradually annexing the translational pattern. All phase patterns converging to the rotational pattern and the translational pattern loses its stability Fig. 3(e), becoming the saddle point. In summary, we demonstrated that our method possesses the ability to configure bifurcations and control the depth of attractors by adjusting the eigenvalues.

The framework can be used to model various coupled multi-body physical system and generate observed or expected spatio-temporal activity patterns. In particular, it can reproduce the dominant large-scale wave patterns extracted from human brain fMRI recordings. Fig. 4 illustrates our modeling of the three low frequency spatio-temporal patterns [Fig. 4(a)] of resting state fMRI obtained by Bolt et al [5]. using the complex principal component analysis (CPCA). The wave patterns are based on Human Connectome Project (HCP) of 50 unrelated participants with *L* = 5124 downsampled cortical vertices as nodes. We assumed that these wave patterns that explain a significant variance are generated from slow oscillations and functional coupling of distributed cortex regions. These dominant patterns are believed to originate from the intrinsic modes of the underlying topology of the coupling network. In view of this, we memorized those three wave patterns with Connetivity strength based method. We first analyzed the stability of phase patterns. Assuming *λ*_*P*1_, *λ*_*P*2_, *λ*_*P*3_ are the eigenvalues corresponding to the three patterns, we calculated the the largest Jacobian matrix eigenvalues of those equilibrium point phase patterns under the parameters space of *λ*_*P*1_, *λ*_*P*2_, *λ*_*P*3_ (keep *λ*_*P*1_ + *λ*_*P*2_ + *λ*_*P*3_ = 1 for normalization). [Fig. 4(c)] shows that by controlling the parameters *λ*_*P*1_, *λ*_*P*2_, *λ*_*P*3_, the model can simultaneously have one, two, or three stable phase patterns. In the adjacent region centered around *λ*_*P*1_ = 1/3, *λ*_*P*2_ = 1/3, *λ*_*P*3_ = 1/3 (marked as black in Fig. 4(c)), the three CPCA components simultaneously become stable attractors. We chose *λ*_*P*1_ = 0.3333, *λ*_*P*2_ = 0.3331, *λ*_*P*3_ = 0.3336 to let the model could simultaneously memory all the three patterns.

**FIG. 4.**
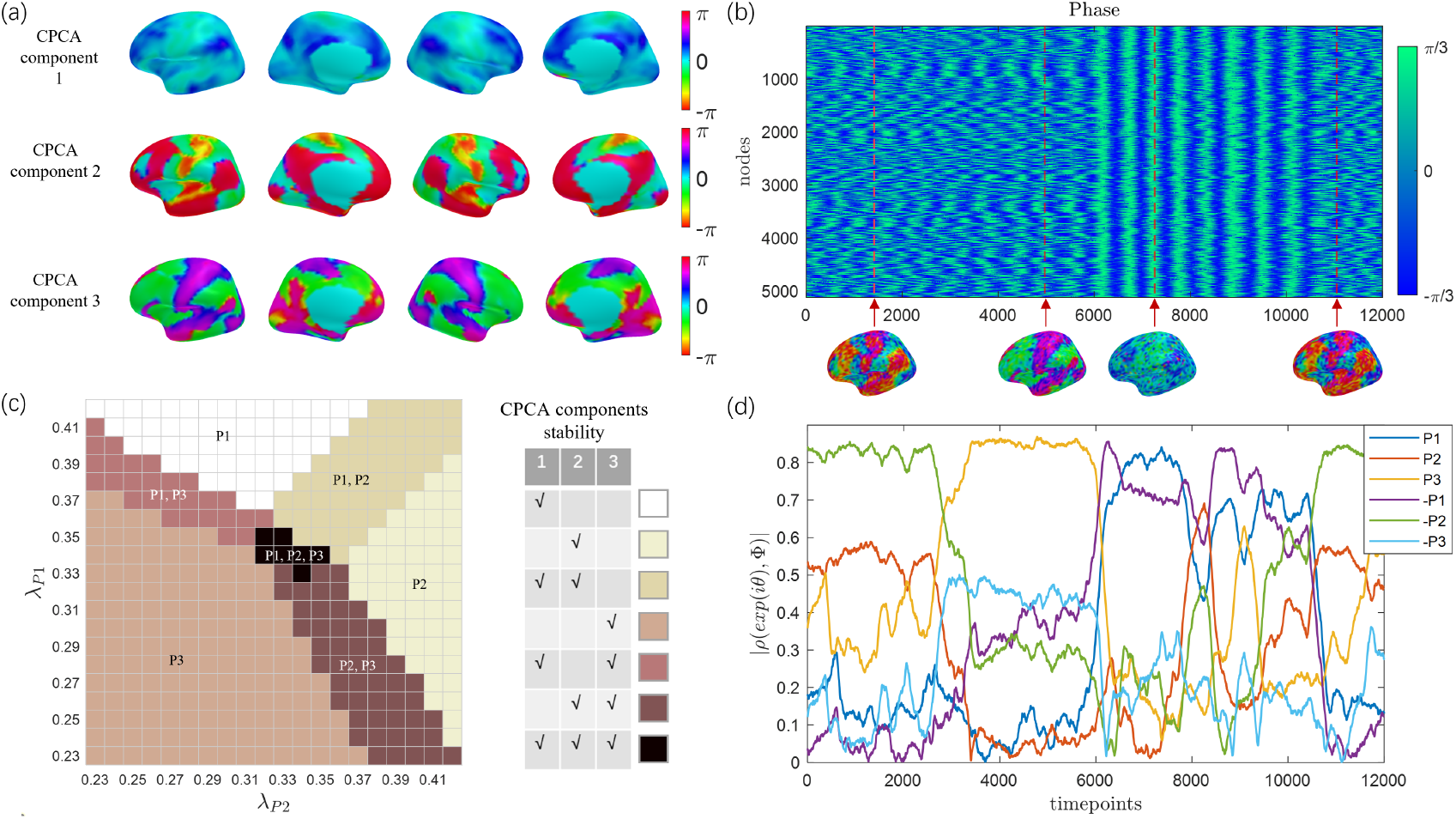
Modeling of dominant spatiotemporal wave patterns in Ref [5]. (a) Three dominant spatio-temporal wave phase patterns in Resting state fMRI captured through complex principal component analysis by Bolt et al [5]. (b) Simulated phase with phase OU noise added, showing 12000 time points captured during the simulation process, including multiple different phase states. The phase patterns at the time points indicated by the arrows (1415, 4969, 7257, 11050) were extracted and presented for demonstration. (c) Stability of three Phase patterns under different eigenvalue parameter *λ*_*P* 1_, *λ*_*P* 2_, *λ*_*P* 3_ configuration (maintained *λ*_*P* 1_ + *λ*_*P* 2_ + *λ*_*P* 3_ = 1), The annotations on the graph indicate the stable phase patterns for different parameter ranges. (d) The absolute value of the Affinity function between simulated phase patterns and three memorized phase patterns (as well as their conjugate phase patterns).

Then, A kind of stochastic kuramoto is introduced in Eq. (20) to reproduce those waves.

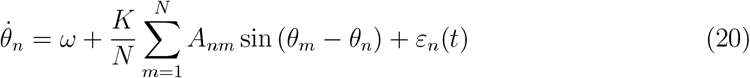

We used Ornstein-Uhlenbeck process (OU) noise *ε*_*n*_(*t*) (see Appendix E for details) to model background noisy activity in the human brain cortex and assume that this noise drives the state transition of wave phase patterns from one attractor to another. We set the local angular frequency for slow oscillation *ω* = 0.1, the global coupling strength *K* = 10 × *L*, set the simulation step to 0.04 s for random initial phase simulation. Meanwhile we calculated the absolute value of the affinity between each time points simulated phase patterns and three memorized patterns (as well as their conjugate phase patterns). [Fig. 4(b, d)] displays the phase states and affinity functions during an intermediate period of the simulation (total duration of 12000 time points, with the starting point of this period denoted as 0). The phase pattern wanders in the attraction domain of 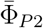 for approximately 2200 time points. Subsequently, the state shifted and Φ_*P*3_ became the dominant mode. Then comes Φ_*P*1_ (Φ_*P*1_ and 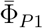 are relatively close). Overall, in most time slots there is a significant dominant wave pattern (∣*ρ* (*e*^*i**θ***(*t*)^, Φ) ∣*>* 0.8), with some periods in unstable transition states. The duration and stability of these states can be adjusted through eigenvalues and input noise parameters.

## IV. CONCLUSION

In this paper, we introduced a framework for elucidating the interplay between spatiotemporal wave patterns and network structure in multi-body systems from the perspective of attractor dynamics. Our approach offers insights into the linkage between the low dimensional neural manifolds and the neural circuits. We derived the characteristic equation for a typical phase-locked state of the Kuramoto coupled oscillators dynamic model. Based on this, we proposed a Hebbian-like heterogeneous connectivity network construction method. It allows us to use attractor dynamics to generate multiple wave phase patterns, as well as control the stability and dominance of each pattern. With this method, We successfully modeled fMRI dominated wave patterns and simulated their dynamic processes, demonstrating a possible forms of fMRI wave spatiotemporal dynamics.

Beside the inter-node phase delays [20], the connectivity strength based method proposed in this study aims to elucidate that the heterogeneous network connectivity can also lead to the specific phase spatial distributions. While the connectivity delay based method generates waves with a preferred directions, the connectivity strength based method allows two opposing waves (conjugate phase patterns) to concurrently serve as attractors. Intriguingly, in the epileptic networks there could exhibits a propagation sequences inversion of upstream and downstream preferences [33], as well as the coexistence of opposing propagation patterns [34]. The electrode microgrids over the human hippocampal surface recorded bidirectional low-frequency oscillations propagation [35]. These phenomena, resembling the model predictions, foreshadow the the presence and influence of heterogeneous connectivity networks.

Our framework presents a significant application prospect in simulating brain activity patterns changing caused by brain state transitions [13]. By manipulating the eigenvalues of brain connectivity networks to control the global dynamic landscape and adding external stimuli or internal perturbations into the whole-brain model, our framework allows for a systematic analysis of the evolution of brain states [36]. It can also inspire the manipulating of these brain wave patterns from the perspective of Hebbian rule neural plasticity. Previous studies have shown that hebbian plasticity can lead to increases in a myelin marker within the fiber bundle [37]. In summary, this study provides a reference mathematical framework for modeling specific spatio-temporal patterns in multi-body systems such as whole brain dynamics.

An open-source code repository for this work is available on GitHub [38].

## Appendix A Phase Pattern Characteristic Equation

We used the mean field method in the complex domain and define that for a *L* nodes kuramoto system, the mean field 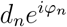 acting on node *n* satisfies

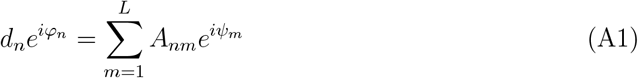

Multiply both sides of the equation by 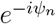 yields

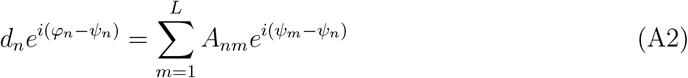

Consider the equality of imaginary parts on both sides of Eq. (A2)

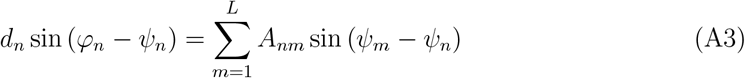

Substituting Eq. (A3) into Eq. (5) we will have *n* differential equations with *ψ*_*n*_ as a variable.

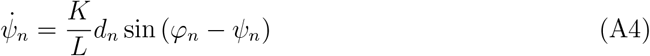

The phase dynamics equations in the mean field sense are completely equivalent to the original differential equations of the equation, and achieve the formally decoupling of multivariable differential equations Eq. (8). To investigate the equilibrium points of phase biased dynamical systems, based on the equilibrium point conditions 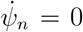, we have *φ*_*n*_ = *ψ*_*n*_. Substituting into Eq. (A1) yields

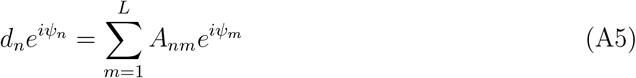

## Appendix B Kuramoto model Jacobian Matrix

For system Eq. (5), we derived its Jacobian matrix ***J***(***ψ***) = {*J*_*nm*_(***ψ***)}_*L*×*L*_ at phase state *ψ*.

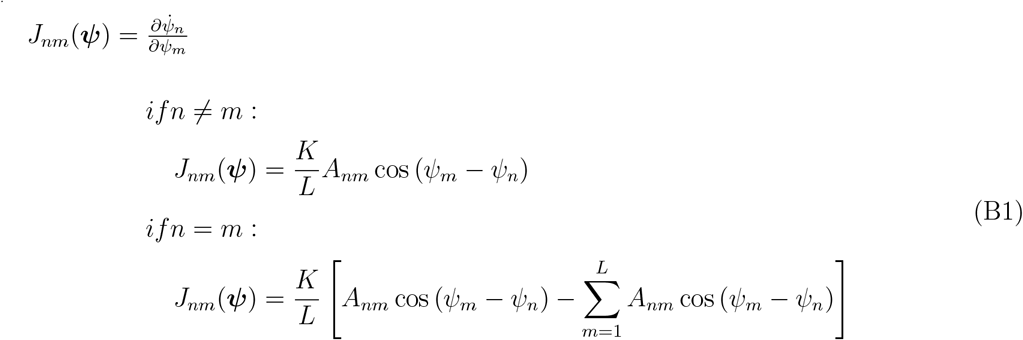

We let ***Q***(***ψ***) = {*Q*_*nm*_(***ψ***)}_*L*×1_ where 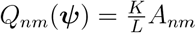 cos (*ψ*_*m*_ − *ψ*_*n*_), It is readily apparent that ***J*** (***ψ***) is the the laplacian matrix of ***Q***(***ψ***).

## Appendix C A General Connectivity Matrix Solution Family

Here we give a more general solution family of connectivity matrices satisfying the Eq. (14)

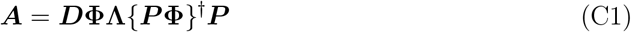

Where ***D*** ∈ ℝ^*L*×*L*^ is a real positive diagonal matrix, ***P*** ∈ ℝ^*L*×*L*^ is a real invertible matrix and other variable symbols are consistent with Eq. (15)

Proof.

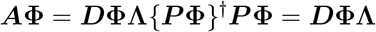

so

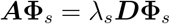

## Appendix D Potential Function and Square Affinity Function

*Proposition*. Let 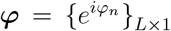 be the pattern memorized via Connectivity strength based method. And ***φ*** satisfies Re(***φ***)^*T*^ Im(***φ***) = 0 (which means ***φ*** is a pure traveling wave) and ∥ Re(***φ***)∥ = ∥ Im(***φ***)∥, where “∥ · ∥” denotes the L2 norm on ℝ^*L*^. Then Eq. (18) holds.

Proof. Consider there are

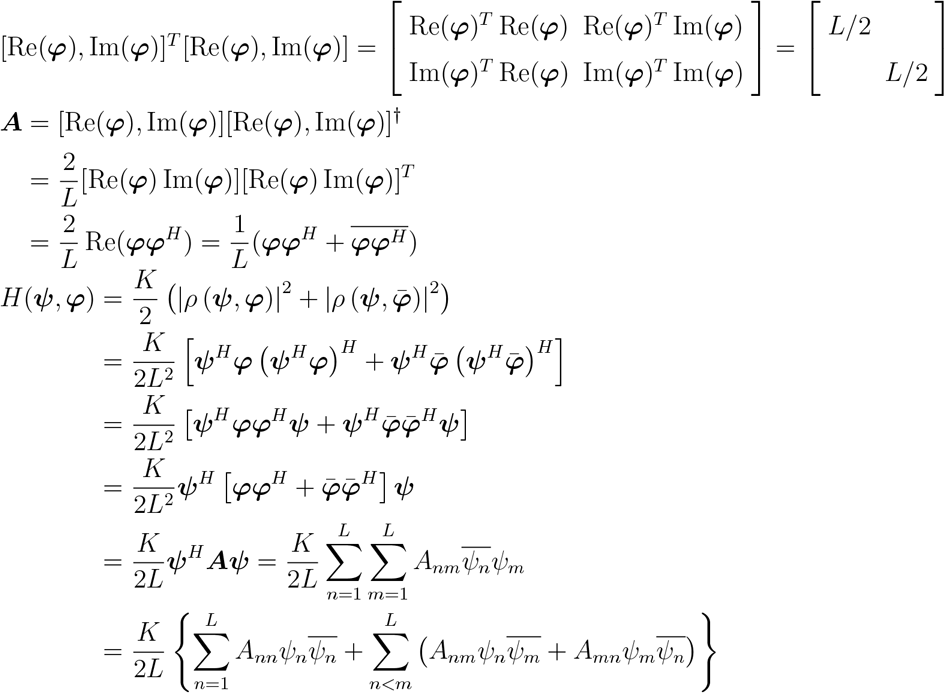

Note that ***A*** is a real symmetric matrix, there are

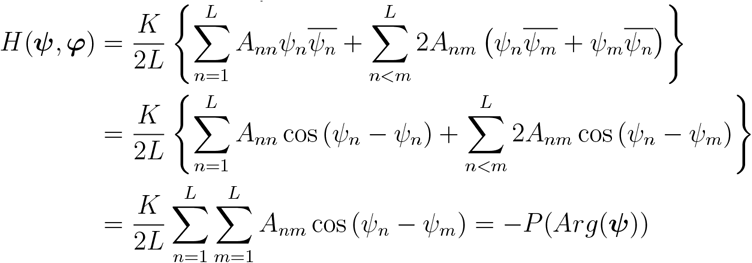

## Appendix E Ornstein-Uhlenbeck Noise

We used the Ornstein-Uhlenbeck stochastic process defined by the following equation to generate OU noise *ε*_*n*_(*t*) for node *n*.

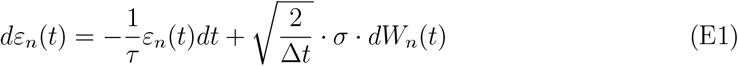

Where *W*_*n*_(*t*) denotes the standard Brownian Motion and independent between nodes. We set *τ* = 300, *σ* = 0.08, and the step size for iteration Δ*t* = 0.08. The generated OU noise from Eq. (E1) is downsampled by a factor of ten for being applied to Eq. (20).

## Notes

### Competing Interest Statement

The authors have declared no competing interest.

### Summary of Updates

We have revised the title of the article and improved the proof process.

